# Baseline Resting-State Functional Connectivity Determines Subsequent Pain Ratings to a Tonic Ecologically Valid Experimental Model of Orofacial Pain

**DOI:** 10.1101/2020.04.28.066555

**Authors:** Lizbeth J. Ayoub, Mary Pat McAndrews, Alexander J. Barnett, Ka Chun Jeremy Ho, Iacopo Cioffi, Massieh Moayedi

## Abstract

Pain is a subjective experience with significant individual differences. Laboratory studies investigating pain thresholds and experimental acute pain have identified structural and functional neural correlates. However, these types of pain stimuli have limited ecological validity to real-life pain experiences. Here, we use an orthodontic procedure—the insertion of an elastomeric separator between teeth—which typically induces mild to moderate pain that peaks within 2 days and lasts several days. We aimed to determine whether the baseline structure and resting-state functional connectivity (rsFC) of key regions along the trigeminal nociceptive and pain modulatory pathways correlate with subsequent peak pain ratings. Twenty-six healthy individuals underwent structural and resting-state functional (rs-fMRI) scanning prior to the placement of a separator between the first and second molars, which was kept in place for five days. Participants recorded pain ratings three times daily on a 100-mm visual analogue scale. Peak pain was not significantly correlated with diffusion metrics of the trigeminal nerve, or grey matter volume of any brain region. Peak pain did, however, positively correlate with baseline rsFC between the thalamus contralateral to the separator and bilateral insula, and negatively correlated with connectivity between the periaqueductal gray (PAG) and core nodes of the default mode network (medial prefrontal and posterior cingulate cortices). The ascending (thalamic) nociceptive and the descending (PAG) pain modulatory pathways at baseline each explained unique variation in peak pain intensity ratings. In sum, pre-interventional functional neural architecture of both systems determined the individual pain experience to a subsequent ecologically valid pain stimulus.

## INTRODUCTION

Pain perception can vary dramatically between individuals: what feels like a light noxious stimulus to one person may feel excruciating to someone else. This experience is shaped by various factors including biological (e.g. genetics), psychological (e.g. anxiety, catastrophizing) and sociocultural factors (e.g. socioeconomic status) [22,23,49,64,66,76,82]. Greater insight into these individual differences has been provided by structural and functional brain imaging studies [23,31,40,43,59-61]. Notably, structural grey matter differences [30,31] and resting-state functional connectivity (rsFC) [21,84] are related to individual differences in pain sensitivity in laboratory-based studies. However, laboratory-based studies of experimental pain are limited in their ecological validity in evaluating a realistic painful experience. Here, we investigate whether pre-interventional (baseline) brain structure and function is related to subsequent pain induced by an ecologically valid and clinically relevant tonic orofacial pain model— orthodontic elastomeric separator insertion.

Orthodontic treatments cause pain in 72-95% of individuals [7,46]. The placement of an elastomeric separator between molars is an orthodontic procedure that creates space between teeth before brace bonding [69]. Orofacial pain induced by the separators in healthy adolescents and adults peaks within 48 hours after insertion, then resolves within 5-7 days [1,13,22]. The separator compresses the periodontal ligament of the alveolar bone, which induces pressure, inflammation and tooth pain [47]. Pain induced by the separator is mediated by the trigeminal nociceptive system and shaped by cognitive-affective factors such as somatosensory amplification and trait anxiety [22]. The trigeminal nerve carries orofacial nociceptive signals to the brainstem [78], and further to the thalamus [15,86] and cortex for processing [29]. The thalamus is the primary relay site of nociceptive input, including from the orofacial region via the trigeminothalamic tract, to the brain [4,15,19,24,27,39,56,70,86,87], and is consistently activated in response to an orofacial noxious stimulus [10]. The experience of pain is modulated by descending circuits. Notably, the periaqueductal grey (PAG), an opiate-rich region, is a key node of these descending circuits which receives convergent input from cortical and subcortical regions, and transmits to the rostral ventral medulla, and further to the spinal cord [35,38,41,54,53,55,57,79,85]. However, whether the structure or functional connectivity of the thalamus and PAG are associated with future individual pain experiences induced by an ecologically valid orofacial pain model has not been investigated.

Here, we aim to determine whether key structures of the ascending trigeminal nociceptive (thalamus) and descending pain modulatory pathways (PAG) relate with peak pain intensity induced by an orthodontic separator placement. We hypothesize that pre-existing structure and functional connectivity of key regions will correlate with an individual’s pain severity in response to this pain model. Specifically, we expect that the structural integrity of the trigeminal nerve and grey matter volume of nociceptive processing and pain modulatory brain regions at baseline will correlate with subsequent peak pain intensity. We further expect that stronger baseline thalamic and weaker PAG functional connectivity with pain-related brain regions will be associated with greater subsequent peak pain intensity. Finally, we expect that each system will contribute uniquely to explaining individual differences in peak pain ratings.

## MATERIALS AND METHODS

### Participants

Twenty-seven healthy individuals consented to a study approval by University of Toronto Human Research Ethics Board (#32797). One participant was unable to complete the study as the separator broke before the end of the five-day experiment, and thus is not included in the analysis. The final cohort comprised 26 participants (11 women, 15 men; aged 25.7 ± 4.4 years). Given that the placement of orthodontic separators is a dental procedure, in Canada, a medical screening by a licensed Orthodontist (IC) is required. As such, participants were screened, and excluded based on the following criteria: current pain, history of chronic pain, chronic illness, psychiatric disorder, pregnancy, metal implants, fixed orthodontic retainers, porcelain-fused-to-metal or full metal crowns, severe dental malocclusions, spaces in the fourth quadrant (lower right mandible; absence of interproximal tooth contacts), undergoing current orthodontic treatment, use of habitual analgesic medication, and presence of dentures. A clinical examination was performed to ensure they did not have temporomandibular disorders (TMD) using the Diagnostic Criteria for Temporomandibular Disorder (DC/TMD) [77]. Participants were financially compensated for their time.

### Pain Model

An elastomeric separator (American Orthodontics, X-Ring Separators, Sheboygan, USA) was placed at the mesial interproximal contact of the right permanent mandibular first molar (fourth quadrant) by a licensed orthodontist (IC). The separator was kept in place for five days, as previously done [22]. Participants were instructed not to take any analgesics, such as ibuprofen or acetaminophen to reduce pain.

### Questionnaires

#### Pain Diary

We provided a pain diary to participants to record their pain intensity three times per day (10:00, 16:00, 22:00) on a 100 mm visual analogue scale (VAS) over the course of five days following separator placement, as previously done [22]. Specifically, participants reported their tooth pain, in response to the following question: “How severe is your tooth pain now? Place a vertical mark on the line below to indicate how bad you feel your tooth pain is now.” The VAS scale had two anchors: left anchor “no pain” and right anchor “worst pain imaginable” [68].

#### Cognitive affective measures

Given that we have previously shown that trait anxiety and somatosensory amplification affected subsequent pain experience induced by the insertion of orthodontic separators [22], we asked participants to complete the following questionnaires before the procedure.

The state-trait anxiety inventory (STAI)-trait anxiety (Y2) is a twenty-item questionnaire, with the scale for each item ranging from 0 to 4 points [83]. It is a self-report questionnaire designed to evaluate trait levels of anxiety. The STAI-state anxiety (Y1) was not used.

The somatosensory amplification scale (SSAS) questionnaire is a ten-item questionnaire, scale for each item ranging from 1 to 5 points [11]. The self-report questionnaire detects whether an individual has a tendency to amplify uncomfortable somatic or visceral sensations.

### Pain rating analyses

To determine pain ratings from the pain diaries, we measured each marking on the pain VAS scale in millimetres and took the average of the three daily pain ratings. As such, each participant had one pain rating per day. The highest average daily pain rating was considered peak pain intensity score for each individual. These pain ratings were used in all neuroimaging analyses.

Daily pain intensity ratings (five per individual) were tested for normality using Shapiro-Wilk’s test. Given that these were not normally distributed, differences between daily ratings were assessed using Friedman’s test, a non-parametric repeated measures ANOVA. Significance was set at *P* < 0.05. Multiple correction of post-hoc tests was performed using Dunn’s test. All statistical analyses on pain ratings were performed using GraphPad v8.4 (https://graphpad.com).

### Relationship between psychological traits and peak pain ratings

Given our previous findings that STAI-trait anxiety and SSAS scores affected the experience of pain induced by elastomeric separators [22], we correlated these psychological measures with peak pain intensity ratings. We tested the STAI-trait and SSAS scores for normality using Shapiro-Wilk’s test. In cases where the data were not normally distributed, we used Spearman’s correlation test to measure the degree of association between each of these scores and peak pain ratings over five days. Significance was set at *P* < 0.05.

### Structural and Functional Neuroimaging

All participants underwent magnetic resonance imaging (MRI) before the insertion of the separator. All scans were acquired using a 3T Siemens Prisma-fit MRI scanner equipped with a 32-channel head coil at the Hospital for Sick Children in Toronto, Canada. Participants were asked to “relax and fixate on the crosshair at the center of the display in the scanner, and not to think about anything in particular.”

#### Structural imaging scan

Structural T1-weighted MRI scans were acquired with a magnetization prepared rapid gradient echo (MPRAGE) using the following sequence: echo time (TE) = 2.96ms; repetition time (TR) = 2300ms; inversion time (TI) = 900ms; 256 sagittal slices; flip angle = 9°; in-plane matrix resolution = 256 × 256 and field-of-view = 256 x 256 mm, resulting in a voxel size = 1 x 1 x 1 mm^3^; with a GRAPPA acceleration factor = 2.

#### Functional imaging scan

Rs-fMRI scans were collected using T2*-weighted echo-planar pulse imaging (EPI) sequence, with the following parameters: TE = 30ms, TR = 1500ms, 50 axial slices; flip angle = 70°; in-plane matrix resolution = 74 × 74 and field-of-view = 222 x 222 mm, resulting in a voxel size = 3 x 3 x 3 mm^3^; 200 volumes, with a multiband slice acceleration factor = 2, total scan time = 5 minutes.

#### Diffusion weighted imaging scan

Two sets of diffusion-weighted images were acquired with reverse phase-encode blips (anterior-posterior and posterior-anterior) resulting in images with distortions in opposite directions with the following parameters: 60 non-collinear diffusion encoding directions with *b* = 1000 s/mm2, 7 non-diffusion encoding volumes (B0). DWI acquisition used the following sequence: TE = 73ms, TR = 4400ms; 80 slices; flip angle = 90°; in-plane matrix resolution = 122 × 122, field-of-view = 244 x 244 mm, resulting in a 2 mm isotropic voxel size; with a multiband slice acceleration factor = 2.

### Neuroimaging analysis

#### Voxel-Based Morphometry

We sought to determine whether brain grey matter volume correlated with peak pain ratings. To do so, we performed a voxel-based morphometry (VBM) analysis on all anatomical T1 scans [6]. VBM was performed in SPM v12 (http://www.fil.ion.ucl.ac.uk/spm/software/spm12) running on MATLAB (R2016b v.9.1; Mathworks, Nantick, MA). First, all scans were centered at the anterior commissure, and were segmented into gray matter, white matter and CSF tissues. Next, the DARTEL toolbox was used to iteratively align grey and white matter images to a study specific template aligned to the MNI template space [5]. Images were smoothed using an 8-mm at full-width at half-maximum (FWHM) Gaussian kernel. We performed a voxelwise general linear model to determine whether grey matter volume correlates with peak pain intensity. Total intracranial volume was included in the general linear model as a nuisance covariate. Significance was set at cluster-corrected *P*_*FDR*_ < 0.05 (with a cluster-forming height threshold of *P* < 0.001). Furthermore, a secondary analysis was run based on rsFC results, restricted to voxels within the seeds and rsFC resulting clusters (see below), significant at cluster-corrected *P*_*FDR*_ < 0.05 (with a cluster-forming height threshold of *P* < 0.001).

#### Diffusion Tensor Imaging

We sought to determine whether microstructure of the cisternal segment of the trigeminal nerve, just outside the pontine trigeminal root-entry zone, is correlated to peak pain ratings. To do so, we performed a diffusion tensor imaging (DTI) analysis using FDT toolbox (FMRIB’s Diffusion Toolbox) in FSL 5.0.11 (https://fsl.fmrib.ox.ac.uk/fsl/fslwiki/FDT). Volumes with no diffusion weighting (B0 volumes) underwent topup, which estimates the susceptibility-induced off-resonance field, i.e. distortion in the subject’s head [2,81]. Next, we used the eddy tool to correct for eddy current distortions, susceptibility-induced distortion, and subject movement [3]. We then fit a tensor model using DTIFIT on the eddy corrected scans. We identified the cisternal segment of the trigeminal nerve, as previously done [20,60], by overlaying the principal eigenvector (V1) map, colored in RGB, onto and modulated to the FA map for each subject in FSLeyes (https://git.fmrib.ox.ac.uk/fsl/fsleyes/fsleyes/). Individual masks of the right cisternal segment of the trigeminal nerve were used to extract DTI metrics, such as fractional anisotropy (FA), mean diffusivity (MD), axial diffusivity (AD) and radial diffusivity (RD) [59]. These measures are calculated from the eigenvalues of the diffusion tensor. FA corresponds to the degree of anisotropic diffusion, ranging from 0 (isotropic diffusion) to 1 (anisotropic diffusion). MD is the average measurement of the three diffusion directions (first, second and third lambdas). AD is the measurement of diffusion along the primary diffusion direction (first lambda). RD represents the average measurement of the secondary and tertiary directions (second and tertiary lambdas). These DTI metrics are indices of microstructural integrity of the nerve. We tested all DTI metrics for normality using Shapiro-Wilk’s test. Then, we tested whether these values were correlated with peak pain ratings using Spearman’s correlation. Significance was set at *P* < 0.05.

#### Seed-to-voxel rsFC analysis

We performed a seed-to-voxel rsFC analysis using CONN v17.f toolbox (http://www.conn-toolbox.org) running on MATLAB (R2016b v.9.1; Mathworks, Nantick, MA). We preprocessed anatomical T1 scans and rs-fMRI scans using tools from the Statistical Parametric Mapping software package (SPM v.12; http://www.fil.ion.ucl.ac.uk/spm/software/spm12) incorporated within CONN. First, we removed five initial scans to allow for field homogeneity. All functional scans underwent realignment, which estimates six parameters of motion and corrects it in the X, Y and Z planes as well as their rotations, roll, pitch, yaw. Next, we used the ART toolbox, implemented in CONN, to detect outlier scans based on conservative settings (global signal threshold at Z=3; subject motion threshold at 0.5 mm) [67]. Following this step, structural and functional scans underwent grey, white and cerebrospinal fluid (CSF) segmentation, and were normalized to the Montreal Neurological Institute (MNI) standard space template. Scans were resliced using default Tissue Probability Maps (structural and functional target resolution at 2 mm). Functional scans were smoothed using an 8-mm FWHM Gaussian kernel. Preprocessed scans underwent denoising, where aCompCor regressed blood-oxygen-level-dependent (BOLD) signal of non-neuronal origin [12], and five principal components were derived for each tissue type (white matter and CSF). Additionally, the following confounding variables were regressed from the model using the indicated number of vectors: white matter (5), CSF (5), motion parameters and their temporal derivatives (12), scrubbing (22) and a regressor to model the first 15 frames and its temporal derivative. This last regressor was added to baseline correct after denoising as normally done in CONN [89]. To identify low-frequency fluctuations characterizing resting state connectivity, we applied a band-pass filter of 0.008-0.09Hz to the data. Then, we performed a whole brain seed-to-voxel rsFC with two seeds: the left thalamus, contralateral to the site of separator placement, and the PAG. The left thalamus region-of-interest (ROI) was anatomically defined based on the FSL Harvard-Oxford atlas, included within CONN. Given the absence of a PAG ROI in any standard neuroimaging atlas, and because of the significant functional heterogeneity of this region, we defined the PAG ROI based on a structural and functional localizer. This approach would selectively include regions of the PAG involved in pain. First, we traced the anatomical borders of the PAG guided by the Duvernoy Atlas [62]. Second, we performed a meta-analytic search on Neurosynth (https://neurosynth.org) with the term “pain”. We extracted the map at a threshold of Z = 17.5 of all the brain regions that showed activation and selected the PAG cluster (86 voxels with a centre-of-gravity at MNI (X, Y, Z): (−1.6, −27.7, −9.88). The overlap between the functional localizer and the structural seed was used in our analysis (Fig S1). For the first-level rsFC analysis, we used bivariate correlations to quantify connectivity between each seed to every other voxel in the brain within each subject. At second-level analysis, we evaluated the effect of peak pain intensity, accounting for sex in the model, using a non-parametric cluster-mass *P*_FDR_ < 0.05 (cluster-mass-forming height threshold *P*_uncorrected_ < 0.001, 1000 permutations). Sex was included in the model as evidence suggests sex differences in pain and in brain networks [32,51,88].

#### Partial correlation analysis between the thalamic and PAG rsFCs and peak pain ratings

We performed a non-parametric partial correlation analysis to determine the ascending nociceptive (thalamic rsFC) and descending pain modulatory pathways’ (PAG rsFC) unique contributions to peak pain ratings using IBM SPSS Statistics for Macintosh v.26 (IBM Corp., Armonk, N.Y., USA). Importantly, the intent of this analysis was not to determine overall correlation values, as that had been achieved in the brain imaging analysis, but rather to identify whether each pathway (the ascending nociceptive pathway and the descending modulatory pathway, captured by connectivity of the thalamus and PAG, respectively) explained unique variance in the pain ratings when accounting for the other’s contribution. Given that suprathreshold clusters were different in voxel number, we exported one mask encompassing the suprathreshold clusters for each seed (one bilateral insular mask of 842 voxels and one mask of 1215 voxels encompassing the right medial prefrontal cortex (mPFC) and the right postcingulate cortex (PCC) in CONN). Then, we used each mask to extract the average rsFC connectivity values. In SPSS, we performed a non-parametric partial correlation between the mean thalamic-insular rsFC connectivity values and peak pain ratings, controlling for the mean PAG-DMN rsFC connectivity values. Conversely, we also performed a non-parametric partial correlation between the mean PAG-DMN rsFC connectivity values and peak pain ratings, controlling for the mean thalamic-insular rsFC connectivity values. Significance for the partial correlation was set at *P* < 0.025, to correct for multiple comparisons from the 2 partial correlations performed.

## RESULTS

### Pain Ratings

Participants rated a median peak pain intensity following the placement of the separator at 17.2 [5.08-31.83] [IQR] mm. Pain ratings were the highest within two days, then declining after the second day (Fig. 1, see Fig. S2 for individual pain ratings). Pain ratings were significantly different across days (χ^2^ = 12.51, *P* = 0.014). Post-hoc paired t-tests revealed that pain ratings on Day 2 (median [IQR] = 8.5 [1.0-25.50] mm) were significantly greater than pain ratings on Day 5 (median [IQR] = 3.3 [1.0-12.42] mm; Dunn-corrected *P* = 0.044).

**Fig. 1.**
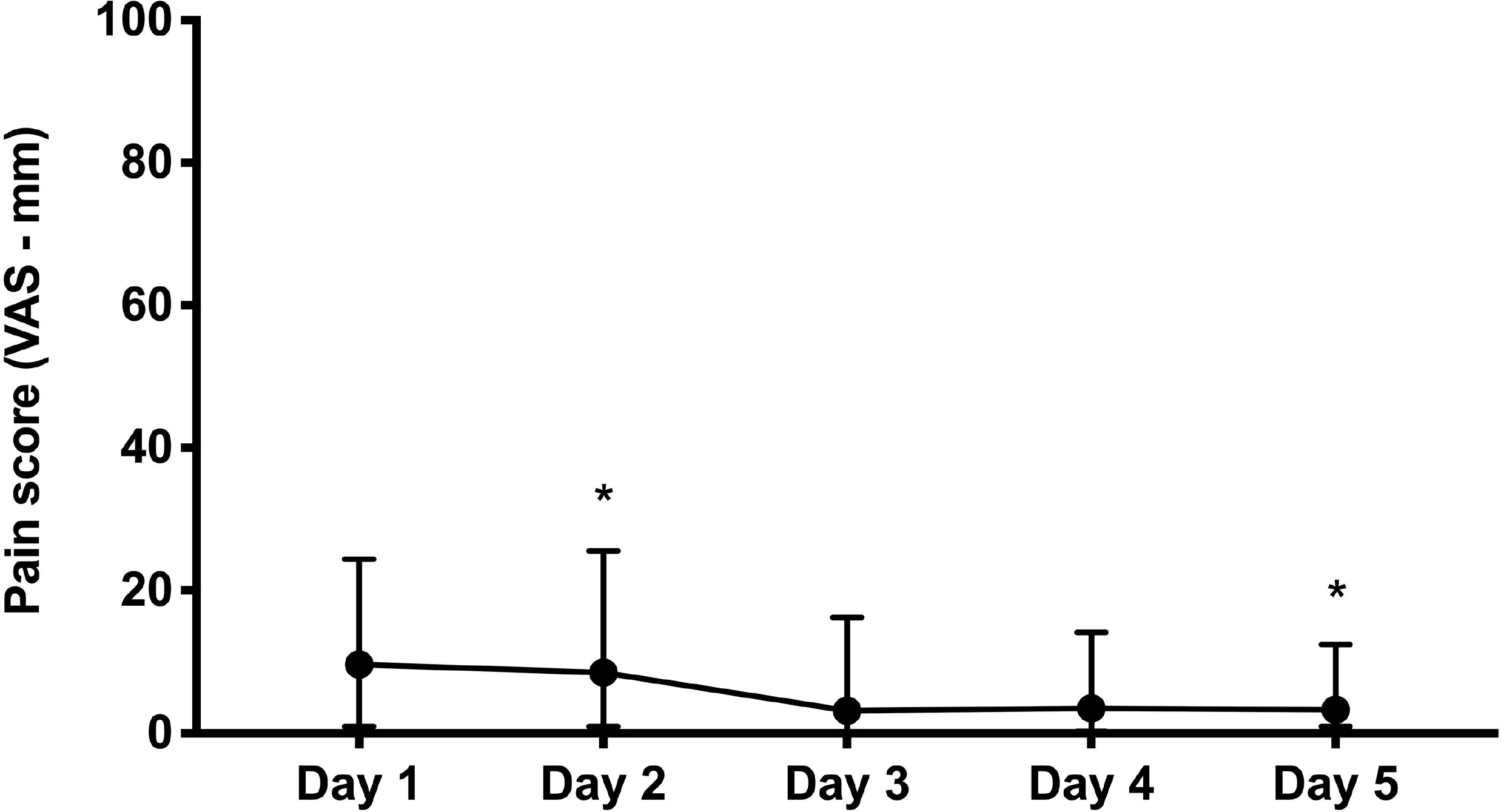
Pain ratings in response to the orthodontic separator for five days. Self-report median pain ratings (IQR) are represented for all participants (n=26) per day. *Pain ratings on Day 2 were significantly increased compared to Day 5, significant at *P* < 0.05. *Abbreviations: IQR* interquartile range *VAS* visual analogue scale.

### Psychological ratings

Participants rated mild trait anxiety (38.3 ± 11.37 (mean ± standard deviation (SD)), and mid-level somatosensory amplification score (16.3 ± 5.76) (mean ± SD). Peak pain ratings were not correlated with trait anxiety (r = 0.12, *P* = 0.55) nor somatosensory amplification scores (r = 0.21, *P* = 0.31).

### Neuroimaging

#### No statistically significant structural correlates of peak pain intensity

We did not find any significant structural correlates of peak pain intensity in our cohort. Specifically, no brain grey matter region’s volume significantly correlated with peak pain intensity. Furthermore, DTI metrics of the right cisternal segment of the trigeminal nerve (mean ± SD: FA = 0.38 ± 0.095; MD = 0.0019 ± 0.00035; RD = 0.0015 ± 0.00037; AD = 0.0027 ± 0.00031) did not significantly correlate with peak pain intensity (FA: r = 0.004, *P* = 0.98; MD: r = 0.004, *P =* 0.98; RD: r = 0.022, *P =* 0.92; and AD: r = −0.031, *P =* 0.88).

#### Stronger thalamo-insular rsFC correlates with peak pain intensity

We found that rsFC between the left thalamus and bilateral insula significantly correlated with subsequent peak pain: left thalamus-left insula (peak MNI coordinates (X,Y,Z) = −36, −8, 0; 597 voxels; 4,776 mm^3^) and left thalamus-right insula (peak MNI coordinates (X,Y,Z) = 40, 6, −6; 245 voxels; 1,960 mm^3^), cluster-mass *P*_FDR_ < 0.05 (Fig. 2). In other words, participants who had stronger baseline connectivity between these regions rated subsequent peak pain as higher.

**Fig. 2.**
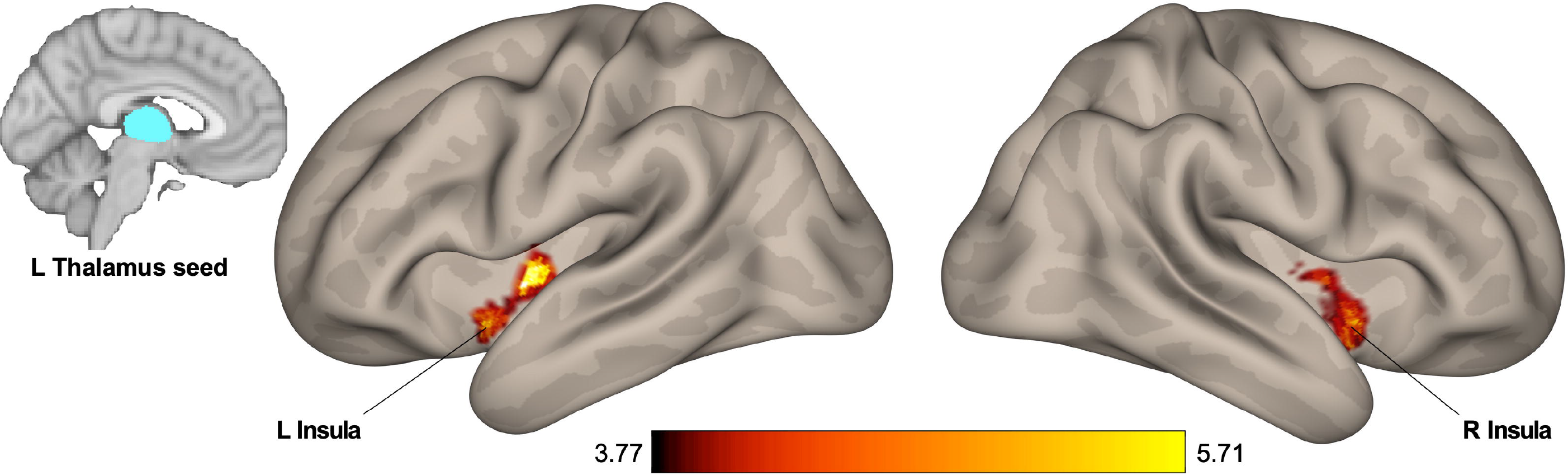
Increased Thalamo-insular rsFC correlates with peak pain intensity. Baseline resting-state functional connectivity (rsFC) between the left thalamus and bilateral insula correlates with subsequent peak pain intensity, significant at cluster-mass *P*_FDR_ < 0.05. Color bar represents T values.

#### Weaker PAG-default mode network rsFC correlates with peak pain intensity

We found that PAG rsFC to nodes of default mode network (DMN) significantly correlated negatively with peak pain intensity: the right mPFC (peak MNI coordinates (X,Y,Z): 12, 44, −12; cluster size: 804 voxels; 6,432 mm^3^) and the right PCC (peak MNI coordinates (X,Y,Z): −8, −48, 14; cluster size: 411 voxels; 3,288 mm^3^), cluster-mass *P*_FDR_ < 0.05 (Fig. 3). In other words, participants who had weaker baseline connectivity between these regions reported higher levels of subsequent peak pain.

**Fig. 3.**
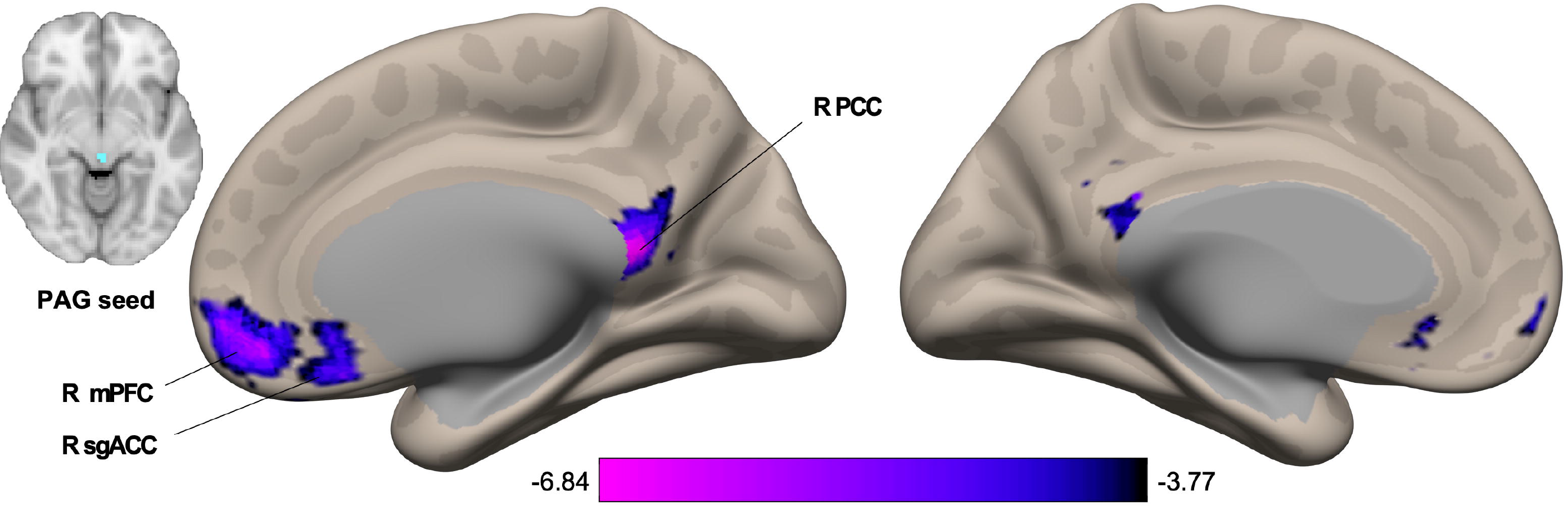
Decreased PAG-DMN correlates with peak pain intensity. Baseline resting-state functional connectivity (rsFC) of the periaqueductal gray (PAG) and core nodes of the default mode network (DMN)—the posterior cingulate cortex (PCC) and medial prefrontal cortex (mPFC) negatively correlates with subsequent peak pain intensity ratings, significant at cluster-mass *P*_FDR_ < 0.05. Color bar represents T values.

#### Ascending nociceptive and descending modulatory systems account for non-overlapping variance of peak pain

To determine whether thalamo-insular connectivity and PAG-DMN connectivity explained non-overlapping (unique) variance in peak pain, we performed two partial correlation analyses. Importantly, connectivity findings were uniquely correlated with peak pain (thalamo-insular connectivity, r = 0.47, P = 0.017, PAG-DMN connectivity, r =-0.63, *P* = 0.001).

## DISCUSSION

Individual variation in pain is thought to be related to the structure and functional connectivity of nociceptive-responsive and pain modulatory regions. Here, we used an ecologically valid model of tonic orofacial pain to evaluate whether baseline structure and rsFC of brain regions involved in nociceptive processing and pain modulation correlate with subsequent peak pain intensity. Pain intensity was highest within 48 hours (Fig. 1). Although we did not find any statistically significant structural correlates of peak pain intensity, we did find that rsFC of key nodes of both the ascending trigeminal nociceptive and descending pain modulatory systems correlated with peak pain. Higher peak pain intensity correlated with stronger thalamo-insular connectivity—key brain regions involved in nociceptive processing (Fig. 2) and weaker rsFC between the PAG with the mPFC and PCC—core nodes of the DMN (Fig. 3). We further showed that thalamo-insular and PAG-DMN connectivities explain unique variance in pain ratings. Together, our data show that individual differences in pre-existing rsFC networks, but not neural structure, can determine the intensity of forthcoming pain in an ecologically valid model of clinical low-level tonic orofacial pain.

Elastomeric separator placement is a common orthodontic procedure. These separators typically induce mild to moderate levels of pain [17]. Our participants had a median of low-level pain compared to previous work in adolescents and adults, who reported moderate levels of pain [1,13,17,22]. However, these studies placed multiple separators between the teeth, which could explain higher pain intensity ratings compared to our study. Since the amount of pain is thought to be proportional to the magnitude of the orthodontic force applied to the teeth [44], increasing the number of separators can be used in future studies to elucidate the effects of different intensities of tonic nociceptive input. Nevertheless, our findings are consistent with these and other studies showing pain intensity is highest within two days of separator placement and resolves by within approximately five days [17,46,63]. Although peak pain intensity in our cohort was variable amongst participants (Fig. S1), the temporal pattern was similar to what has been reported, such that Day 2 had a significantly higher median rating than Day 5.

Separator placement is an ecologically valid orofacial pain model. It circumvents the limitations of laboratory-based pain testing, where stimuli are usually brief (i.e., do not go beyond the experimental session) and only capture a brief snapshot of the experience in a controlled setting [72]. This model captures environmental variability, such that the individual leaves the laboratory setting, while continuously receiving the stimulus for the duration of the separator placement. This ecological validity is fundamental to capturing individual differences in pain, particularly in its evolution and resolution.

Brain structure is thought to reflect, and even predict individual differences in behaviour and cognition [45]. A limited of number of studies in healthy adults have reported baseline gray matter correlates of acute laboratory-based thermal pain stimuli [30,31,34] in various brain regions. In individuals with trigeminal neuralgia, diffusion metrics of the trigeminal nerve have been used to predict treatment responsiveness [43]. However, our study did not identify any significant structural correlates of peak pain, even with a directed search of regions identified in the rsFC analysis. Notably, previous gray matter studies had substantially larger sample sizes and/or small effect sizes. Therefore, our sample size may have been too small to detect subtle effects. For the trigeminal nerve, the FA values in our cohort were within a normative range, whereas those for the affected nerve in trigeminal neuralgia cohorts were abnormally low [43]. Therefore, it is likely that structural white matter abnormalities may predict improvement after treatment; whereas FA variability (and other diffusion metrics) may not be good predictors of nociceptive transmission in the absence of pathology. Therefore, neural structure may not be a suitable correlate for this ecologically valid model of orofacial pain in otherwise healthy individuals.

Brain networks show spontaneous variations in correlated activity of constituents at rest, including those involved in nociceptive processing and pain modulation; they can be thought to reflect neural configurations that can be mobilized for specific types of information processing [80]. Previous studies in healthy adults show increased rsFC between brain regions involved in pain processing before receiving nociceptive stimuli, i.e. pain anticipation [16,65] and following nociceptive stimulation [74]. Analogous to the current findings, a recent study demonstrated that data-driven, multivariate analyses of rsFC in a pain-free state predicted individual differences in pain sensitivity during later testing [84]. Their measure of pain sensitivity was a ‘composite’ based on a weighted average of heat, cold and mechanical pain thresholds delivered 1-3 days from the time of resting-state functional imaging. Notably, pain thresholds measure a brief static point at which the stimulus becomes painful for the individual [75], and do not reflect most real-life painful experiences. In contrast, our study shows that both baseline ascending and descending systems determine subsequent individual peak pain induced by an ecologically valid tonic noxious stimulus.

We investigated the rsFC of the thalamus, a major relay region of the ascending nociceptive pathways [26]. The thalamus and insula are structurally connected. Evidence from primates and humans show efferent and afferent projections from the dorsal thalamus to the insular cortex [9]. The dorsal thalamus projects to the insula from the ventral anterior, centromedian, ventral posterior medial nuclei [8]. Specifically, there is evidence that the ventral medial posterior thalamus projects to posterior insula [25,26,28]. There is a functional dichotomy in the involvement of the insula in pain perception: the anterior insula is closely associated with the cognitive and/or modulatory aspect of pain, whereas the posterior and mid-insula are associated with lower-level, sensory-discriminative features of pain, such as location and intensity [18,58,90]. Some have even proposed that the posterior insula/opercular region could serve as primary pain cortex [33]. Our insular clusters are located in the posterior and mid-insula, and therefore receive direct input from the thalamus, and may reflect the first cortical relays of nociceptive information processing. Therefore, our study advances this finding by showing that baseline thalamo-insular connectivity correlates with subsequent peak pain intensity, and thus supports the sensory-discriminative role of the insula in pain.

Pain perception is a fine balance between nociceptive and antinociceptive mechanisms [14]. Early evidence in rodents [73] and humans [42] implicate the PAG in pain inhibitory mechanisms. More recent evidence suggests that the PAG is involved in both facilitatory and inhibitory pain mechanisms [52]. Our results show that peak pain intensity was negatively correlated with PAG-DMN rsFC. The PAG receives input from ascending spinal/trigeminal systems, descending input from cortical subcortical regions. Further, it projects to brainstem pain modulatory circuits, and is a source of opioidergic pain modulation [36,52,57]. Specifically, the dorsolateral and lateral subregions of the PAG receive input from the mPFC and PCC [37,36,50,52]. The mPFC, PCC and lateral parietal cortices are regions whose activity is highly correlated and form the DMN [71]. DMN activity increases when the individual is not attending to external stimuli and is involved in monitoring the internal milieu which would include monitoring nociceptive signals. We found that individuals who felt less pain had a stronger baseline PAG-DMN rsFC. Previous evidence shows that stronger PAG-DMN rsFC is associated with the tendency to disengage from pain, in other words mind wander away from pain [48]. Thus, high baseline connectivity between the PAG and DMN may enable greater engagement of descending systems in order to disengage from pain. Conversely, weaker PAG-mPFC rsFC in patients with chronic low back pain was related to higher pain intensity conditions, suggesting dysfunction in the descending pain modulation [91]. As such, our study further postulates that the strength of PAG-DMN rsFC at baseline in the healthy individuals correlates with subsequent peak pain intensity. Whether this rsFC could predict either the development or maintenance of chronic pain requires further investigation.

We further demonstrate unique contributions of the ascending nociceptive system and descending pain modulatory pathway to subsequent peak pain ratings. However, our study is limited by our relatively small sample size, and thus we could not further parse the contributions of sex-differences which are known to affect pain ratings in healthy adults. Another limitation is the nature of rsFC, which may differ from brain activity during nociceptive processing, and is thus limited in delineating the neural mechanisms of nociceptive processing. Nonetheless, our findings and others that indicate baseline neural functional architecture is correlated to a future painful experience suggests that these can be leveraged to predict individual pain responses. Future studies with larger sample sizes should test whether these relationships will hold in independent samples and develop predictive models. Understanding a healthy individual’s pain response is an important avenue of research, which could help guide clinical decision making based on an individual’s predicted response to a particular treatment.

In sum, we suggest that the pre-existing state of the ascending and descending nociceptive pathways may determine subsequent pain ratings to an ecologically valid orofacial pain model and that each system contributes uniquely to the individual’s pain response.

## ACKNOWLEDGEMENTS

We would like to thank Ms. Sinéad Devitt for her help with the diffusion imaging analysis. LA is supported by Canadian Institutes of Health Research (CIHR) Frederick Banting and Charles Best Doctoral Research Award. AB is supported by an NSERC Postdoctoral Fellowship Award. KCJH was supported by a CIHR summer studentship award. MM acknowledges support from the Bertha Rosenstadt Endowment Fund. He also holds a Natural Sciences and Engineering Research Council (Canada) Discovery Grant RGPIN-2018-04908. The study was funded by the American Association of Orthodontists Foundation through an Orthodontic Faculty Development Fellowship awarded to IC. The authors declare no competing financial interests.

## AUTHORS CONTRIBUTIONS

LA: data analysis and drafting of manuscript; MPM: revising the manuscript; AB: data analysis; KCJH: data collection; IC and MM: conception and study design, revising the manuscript. All authors have approved the final version and agree to be accountable for all aspects of the work.

## REFERENCES

[1] Aldrees AM. Intensity of pain due to separators in adolescent orthodontic patients. J Orthod Sci 4:118–122, 2015.

[2] Andersson JL, Skare S, Ashburner J. How to correct susceptibility distortions in spin-echo echoplanar images: application to diffusion tensor imaging. Neuroimage 20:870–888, 2003.

[3] Andersson JLR, Sotiropoulos SN. An integrated approach to correction for off-resonance effects and subject movement in diffusion MR imaging. Neuroimage 125:1063–1078, 2016.

[4] Apkarian AV. A brain signature for acute pain. Trends Cogn Sci 17:309–310, 2013.

[5] Ashburner J. A fast diffeomorphic image registration algorithm. Neuroimage 38:95–113, 2007.

[6] Ashburner J, Friston KJ. Voxel-based morphometry--the methods. Neuroimage 11:805–821, 2000.

[7] Asiry MA, Albarakati SF, Al-Marwan MS, Al-Shammari RR. Perception of pain and discomfort from elastomeric separators in Saudi adolescents. Saudi Med J 35:504–507, 2014.

[8] Augustine JR. The insular lobe in primates including humans. Neurol Res 7:2–10, 1985.

[9] Augustine JR. Circuitry and functional aspects of the insular lobe in primates including humans. Brain Res Brain Res Rev 22:229–244, 1996.

[10] Ayoub LJ, Seminowicz DA, Moayedi M. A meta-analytic study of experimental and chronic orofacial pain excluding headache disorders. NeuroImage Clinical 20:901–912, 2018.

[11] Barsky AJ, Goodson JD, Lane RS, Cleary PD. The amplification of somatic symptoms. Psychosom Med 50:510–519, 1988.

[12] Behzadi Y, Restom K, Liau J, Liu TT. A component based noise correction method (CompCor) for BOLD and perfusion based fMRI. Neuroimage 37:90–101, 2007.

[13] Bergius M, Berggren U, Kiliaridis S. Experience of pain during an orthodontic procedure. Eur J Oral Sci 110:92–98, 2002.

[14] Bingel U, Schoell E, Herken W, Buchel C, May A. Habituation to painful stimulation involves the antinociceptive system. Pain 131:21–30, 2007.

[15] Boivie J. An anatomical reinvestigation of the termination of the spinothalamic tract in the monkey. J Comp Neurol 186:343–369, 1979.

[16] Boly M, Balteau E, Schnakers C, Degueldre C, Moonen G, Luxen A, Phillips C, Peigneux P, Maquet P, Laureys S. Baseline brain activity fluctuations predict somatosensory perception in humans. Proc Natl Acad Sci U S A 104:12187–12192, 2007.

[17] Bondemark L, Fredriksson K, Ilros S. Separation effect and perception of pain and discomfort from two types of orthodontic separators. World J Orthod 5:172–176, 2004.

[18] Brooks JC, Zambreanu L, Godinez A, Craig AD, Tracey I. Somatotopic organisation of the human insula to painful heat studied with high resolution functional imaging. Neuroimage 27:201–209, 2005.

[19] Bushnell MC, Duncan GH, Tremblay N. Thalamic VPM nucleus in the behaving monkey. I. Multimodal and discriminative properties of thermosensitive neurons. J Neurophysiol 69:739–752, 1993.

[20] Chen DQ, DeSouza DD, Hayes DJ, Davis KD, O’Connor P, Hodaie M. Diffusivity signatures characterize trigeminal neuralgia associated with multiple sclerosis. Mult Scler 22:51–63, 2016.

[21] Cheng JC, Erpelding N, Kucyi A, DeSouza DD, Davis KD. Individual Differences in Temporal Summation of Pain Reflect Pronociceptive and Antinociceptive Brain Structure and Function. J Neurosci 35:9689–9700, 2015.

[22] Cioffi I, Michelotti A, Perrotta S, Chiodini P, Ohrbach R. Effect of somatosensory amplification and trait anxiety on experimentally induced orthodontic pain. Eur J Oral Sci 124:127–134, 2016.

[23] Coghill RC, McHaffie JG, Yen YF. Neural correlates of interindividual differences in the subjective experience of pain. Proc Natl Acad Sci U S A 100:8538–8542, 2003.

[24] Coghill RC, Sang CN, Maisog JM, Iadarola MJ. Pain intensity processing within the human brain: a bilateral, distributed mechanism. J Neurophysiol 82:1934–1943, 1999.

[25] Craig AD. Topographically organized projection to posterior insular cortex from the posterior portion of the ventral medial nucleus in the long-tailed macaque monkey. J Comp Neurol 522:36–63, 2014.

[26] Craig AD, Bushnell MC, Zhang ET, Blomqvist A. A thalamic nucleus specific for pain and temperature sensation. Nature 372:770–773, 1994.

[27] Davis KD, Dostrovsky JO. Properties of feline thalamic neurons activated by stimulation of the middle meningeal artery and sagittal sinus. Brain Res 454:89–100, 1988.

[28] Davis KD, Lozano RM, Manduch M, Tasker RR, Kiss ZH, Dostrovsky JO. Thalamic relay site for cold perception in humans. J Neurophysiol 81:1970–1973, 1999.

[29] Davis KD, Moayedi M. Central Mechanisms of Pain Revealed Through Functional and Structural MRI. Journal of neuroimmune pharmacology: the official journal of the Society on NeuroImmune Pharmacology 8:518–534, 2013.

[30] Emerson NM, Zeidan F, Lobanov OV, Hadsel MS, Martucci KT, Quevedo AS, Starr CJ, Nahman-Averbuch H, Weissman-Fogel I, Granovsky Y, Yarnitsky D, Coghill RC. Pain sensitivity is inversely related to regional grey matter density in the brain. Pain 155:566–573, 2014.

[31] Erpelding N, Moayedi M, Davis KD. Cortical thickness correlates of pain and temperature sensitivity. Pain 153:1602–1609, 2012.

[32] Galli G, Santarnecchi E, Feurra M, Bonifazi M, Rossi S, Paulus MP, Rossi A. Individual and sex-related differences in pain and relief responsiveness are associated with differences in resting-state functional networks in healthy volunteers. Eur J Neurosci 43:486–493, 2016.

[33] Garcia-Larrea L. The posterior insular-opercular region and the search of a primary cortex for pain. Neurophysiol Clin 42:299–313, 2012.

[34] Grant JA, Courtemanche J, Duerden EG, Duncan GH, Rainville P. Cortical thickness and pain sensitivity in zen meditators. Emotion 10:43–53, 2010.

[35] Hadjipavlou G, Dunckley P, Behrens TE, Tracey I. Determining anatomical connectivities between cortical and brainstem pain processing regions in humans: a diffusion tensor imaging study in healthy controls. Pain 123:169–178, 2006.

[36] Hardy SG, Leichnetz GR. Cortical projections to the periaqueductal gray in the monkey: a retrograde and orthograde horseradish peroxidase study. Neurosci Lett 22:97–101, 1981.

[37] Hardy SG, Leichnetz GR. Frontal cortical projections to the periaqueductal gray in the rat: a retrograde and orthograde horseradish peroxidase study. Neurosci Lett 23:13–17, 1981.

[38] Hayashi H, Sumino R, Sessle BJ. Functional organization of trigeminal subnucleus interpolaris: nociceptive and innocuous afferent inputs, projections to thalamus, cerebellum, and spinal cord, and descending modulation from periaqueductal gray. J Neurophysiol 51:890–905, 1984.

[39] Head H, Holmes G. Sensory disturbances from cerebral lesions. Book Sensory disturbances from cerebral lesions. City: Royal College of Physicians, 1911.

[40] Hemington KS, Cheng JC, Bosma RL, Rogachov A, Kim JA, Davis KD. Beyond Negative Pain-Related Psychological Factors: Resilience Is Related to Lower Pain Affect in Healthy Adults. J Pain 18:1117–1128, 2017.

[41] Hemington KS, Coulombe MA. The periaqueductal gray and descending pain modulation: why should we study them and what role do they play in chronic pain? J Neurophysiol 114:2080–2083, 2015.

[42] Hosobuchi Y, Adams JE, Linchitz R. Pain relief by electrical stimulation of the central gray matter in humans and its reversal by naloxone. Science 197:183–186, 1977.

[43] Hung PS, Chen DQ, Davis KD, Zhong J, Hodaie M. Predicting pain relief: Use of pre-surgical trigeminal nerve diffusion metrics in trigeminal neuralgia. NeuroImage Clinical 15:710–718, 2017.

[44] Jones M, Chan C. The pain and discomfort experienced during orthodontic treatment: a randomized controlled clinical trial of two initial aligning arch wires. Am J Orthod Dentofacial Orthop 102:373–381, 1992.

[45] Kanai R, Rees G. The structural basis of inter-individual differences in human behaviour and cognition. Nat Rev Neurosci 12:231–242, 2011.

[46] Kavaliauskiene A, Smailiene D, Buskiene I, Keriene D. Pain and discomfort perception among patients undergoing orthodontic treatment: results from one month follow-up study. Stomatologija 14:118–125, 2012.

[47] Krishnan V. Orthodontic pain: from causes to management--a review. Eur J Orthod 29:170–179, 2007.

[48] Kucyi A, Salomons TV, Davis KD. Mind wandering away from pain dynamically engages antinociceptive and default mode brain networks. Proc Natl Acad Sci U S A 110:18692–18697, 2013.

[49] Lautenbacher S, Peters JH, Heesen M, Scheel J, Kunz M. Age changes in pain perception: A systematic-review and meta-analysis of age effects on pain and tolerance thresholds. Neurosci Biobehav Rev 75:104–113, 2017.

[50] Leonard CM. The prefrontal cortex of the rat. I. Cortical projection of the mediodorsal nucleus. II. Efferent connections. Brain Res 12:321–343, 1969.

[51] Linnman C, Beucke JC, Jensen KB, Gollub RL, Kong J. Sex similarities and differences in pain-related periaqueductal gray connectivity. Pain 153:444–454, 2012.

[52] Linnman C, Moulton EA, Barmettler G, Becerra L, Borsook D. Neuroimaging of the periaqueductal gray: state of the field. Neuroimage 60:505–522, 2012.

[53] Mantyh PW. Connections of midbrain periaqueductal gray in the monkey. I. Ascending efferent projections. J Neurophysiol 49:567–581, 1983.

[54] Mantyh PW. Connections of midbrain periaqueductal gray in the monkey. II. Descending efferent projections. J Neurophysiol 49:582–594, 1983.

[55] Mantyh PW, Peschanski M. Spinal projections from the periaqueductal grey and dorsal raphe in the rat, cat and monkey. Neuroscience 7:2769–2776, 1982.

[56] Mehler WR, Feferman ME, Nauta WJ. Ascending axon degeneration following anterolateral cordotomy. An experimental study in the monkey. Brain 83:718–750, 1960.

[57] Millan MJ. Descending control of pain. Prog Neurobiol 66:355–474, 2002.

[58] Moayedi M. All roads lead to the insula. Pain 155:1920–1921, 2014.

[59] Moayedi M, Hodaie M. Trigeminal nerve and white matter brain abnormalities in chronic orofacial pain disorders. Pain Rep 4:e755, 2019.

[60] Moayedi M, Weissman-Fogel I, Salomons TV, Crawley AP, Goldberg MB, Freeman BV, Tenenbaum HC, Davis KD. White matter brain and trigeminal nerve abnormalities in temporomandibular disorder. Pain 153:1467–1477, 2012.

[61] Moulton EA, Keaser ML, Gullapalli RP, Maitra R, Greenspan JD. Sex differences in the cerebral BOLD signal response to painful heat stimuli. Am J Physiol Regul Integr Comp Physiol 291:R257–267, 2006.

[62] Naidich T, Duvernoy H, Delman B, Sorensen A, Kollias S, Haacke E. Duvernoy’s atlas: The human brain stem & cerebellum. High-field MRI: Surface anatomy, internal structure, vascularization & 3D sectional anatomy: Springer-Verlag, 2007.

[63] Ngan P, Kess B, Wilson S. Perception of discomfort by patients undergoing orthodontic treatment. Am J Orthod Dentofacial Orthop 96:47–53, 1989.

[64] Ong AD, Zautra AJ, Reid MC. Psychological resilience predicts decreases in pain catastrophizing through positive emotions. Psychol Aging 25:516–523, 2010.

[65] Ploner M, Lee MC, Wiech K, Bingel U, Tracey I. Prestimulus functional connectivity determines pain perception in humans. Proc Natl Acad Sci U S A 107:355–360, 2010.

[66] Poleshuck EL, Green CR. Socioeconomic disadvantage and pain. Pain 136:235–238, 2008.

[67] Power JD, Barnes KA, Snyder AZ, Schlaggar BL, Petersen SE. Spurious but systematic correlations in functional connectivity MRI networks arise from subject motion. Neuroimage 59:2142–2154, 2012.

[68] Price DD, McGrath PA, Rafii A, Buckingham B. The validation of visual analogue scales as ratio scale measures for chronic and experimental pain. Pain 17:45–56, 1983.

[69] Proffit WR, Fields HW, Larson BE, Sarver DM. Contemporary Orthodontics: Elsevier Health Sciences, 2019.

[70] Quensel F. Neurol Zbl 17:482, 1898.

[71] Raichle ME. The brain’s default mode network. Annu Rev Neurosci 38:433–447, 2015.

[72] Reddy KS, Naidu MU, Rani PU, Rao TR. Human experimental pain models: A review of standardized methods in drug development. J Res Med Sci 17:587–595, 2012.

[73] Reynolds DV. Surgery in the rat during electrical analgesia induced by focal brain stimulation. Science 164:444–445, 1969.

[74] Riedl V, Valet M, Woller A, Sorg C, Vogel D, Sprenger T, Boecker H, Wohlschlager AM, Tolle TR. Repeated pain induces adaptations of intrinsic brain activity to reflect past and predict future pain. Neuroimage 57:206–213, 2011.

[75] Rolke R, Magerl W, Campbell KA, Schalber C, Caspari S, Birklein F, Treede RD. Quantitative sensory testing: a comprehensive protocol for clinical trials. Eur J Pain 10:77–88, 2006.

[76] Rollman GB, Abdel-Shaheed J, Gillespie JM, Jones KS. Does past pain influence current pain: biological and psychosocial models of sex differences. Eur J Pain 8:427–433, 2004.

[77] Schiffman E, Ohrbach R, Truelove E, Look J, Anderson G, Goulet JP, List T, Svensson P, Gonzalez Y, Lobbezoo F, Michelotti A, Brooks SL, Ceusters W, Drangsholt M, Ettlin D, Gaul C, Goldberg LJ, Haythornthwaite JA, Hollender L, Jensen R, John MT, De Laat A, de Leeuw R, Maixner W, van der Meulen M, Murray GM, Nixdorf DR, Palla S, Petersson A, Pionchon P, Smith B, Visscher CM, Zakrzewska J, Dworkin SF, International Rdc/Tmd Consortium Network IafDR, Orofacial Pain Special Interest Group IAftSoP. Diagnostic Criteria for Temporomandibular Disorders (DC/TMD) for Clinical and Research Applications: recommendations of the International RDC/TMD Consortium Network* and Orofacial Pain Special Interest Groupdagger. J Oral Facial Pain Headache 28:6–27, 2014.

[78] Sessle BJ. Neural mechanisms and pathways in craniofacial pain. Can J Neurol Sci 26 Suppl 3:S7–11, 1999.

[79] Sillery E, Bittar RG, Robson MD, Behrens TE, Stein J, Aziz TZ, Johansen-Berg H. Connectivity of the human periventricular-periaqueductal gray region. J Neurosurg 103:1030–1034, 2005.

[80] Smith SM, Fox PT, Miller KL, Glahn DC, Fox PM, Mackay CE, Filippini N, Watkins KE, Toro R, Laird AR, Beckmann CF. Correspondence of the brain’s functional architecture during activation and rest. Proc Natl Acad Sci U S A 106:13040–13045, 2009.

[81] Smith SM, Jenkinson M, Woolrich MW, Beckmann CF, Behrens TE, Johansen-Berg H, Bannister PR, De Luca M, Drobnjak I, Flitney DE, Niazy RK, Saunders J, Vickers J, Zhang Y, De Stefano N, Brady JM, Matthews PM. Advances in functional and structural MR image analysis and implementation as FSL. Neuroimage 23 Suppl 1:S208–219, 2004.

[82] Sorge RE, Totsch SK. Sex Differences in Pain. J Neurosci Res 95:1271–1281, 2017.

[83] Spielberger CD, Gorusch RL, Lushene RE. Manual of the State-Trait Anxiety Inventory, 1970.

[84] Spisak T, Kincses B, Schlitt F, Zunhammer M, Schmidt-Wilcke T, Kincses ZT, Bingel U. Pain-free resting-state functional brain connectivity predicts individual pain sensitivity. Nat Commun 11:187, 2020.

[85] Tracey I, Mantyh PW. The cerebral signature for pain perception and its modulation. Neuron 55:377–391, 2007.

[86] Treede RD, Kenshalo DR, Gracely RH, Jones AK. The cortical representation of pain. Pain 79:105–111, 1999.

[87] Vogt BA. Pain and emotion interactions in subregions of the cingulate gyrus. Nat Rev Neurosci 6:533–544, 2005.

[88] Wang G, Erpelding N, Davis KD. Sex differences in connectivity of the subgenual anterior cingulate cortex. Pain 155:755–763, 2014.

[89] Whitfield-Gabrieli S, Nieto-Castanon A. Conn: A functional connectivity toolbox for correlated and anticorrelated brain networks. Brain connectivity 2:125–141, 2012.

[90] Wiech K, Jbabdi S, Lin CS, Andersson J, Tracey I. Differential structural and resting state connectivity between insular subdivisions and other pain-related brain regions. Pain 155:2047–2055, 2014.

[91] Yu R, Gollub RL, Spaeth R, Napadow V, Wasan A, Kong J. Disrupted functional connectivity of the periaqueductal gray in chronic low back pain. NeuroImage Clinical 6:100–108, 2014.

