## Supplementary material for "Baseline Resting-State Functional Connectivity Determines Subsequent Pain Ratings to a Tonic Ecologically Valid Experimental Model of Orofacial Pain": Fig. S

**Abbreviated title:** Baseline rsFC Determines Subsequent Pain Ratings

Lizbeth J. Ayoub^1,2,3^, Mary Pat McAndrews^2,4^, Alexander J. Barnett^5^, Ka Chun Jeremy Ho^1^, Iacopo Cioffi^1,3,6^, Massieh Moayedi^1,3,6§^

^1^Centre for Multimodal Sensorimotor and Pain Research, Faculty of Dentistry, University of Toronto, Toronto, ON, Canada, M5G 1E2;

^2^Division of Clinical and Computational Neuroscience, Krembil Brain Institute, Toronto

Western Hospital, University Health Network, Toronto, ON, Canada, M5T 2S8;

^3^University of Toronto Centre for the Study of Pain, Toronto, ON, Canada, M5T 1P8;

^4^Department of Psychology, University of Toronto, Toronto, ON, Canada, M5S 3G3;

^5^Center for Neuroscience, University of California at Davis, Davis, California, USA;

^6^Department of Dentistry, Mount Sinai Hospital, Toronto, ON, Canada, M5G 1X5.

§Please address all correspondence to:

Massieh Moayedi, PhD

Centre for Multimodal Sensorimotor and Pain Research

Faculty of Dentistry - University of Toronto

123 Edward Street, Room 501B

Toronto, ON

M5G 1E2

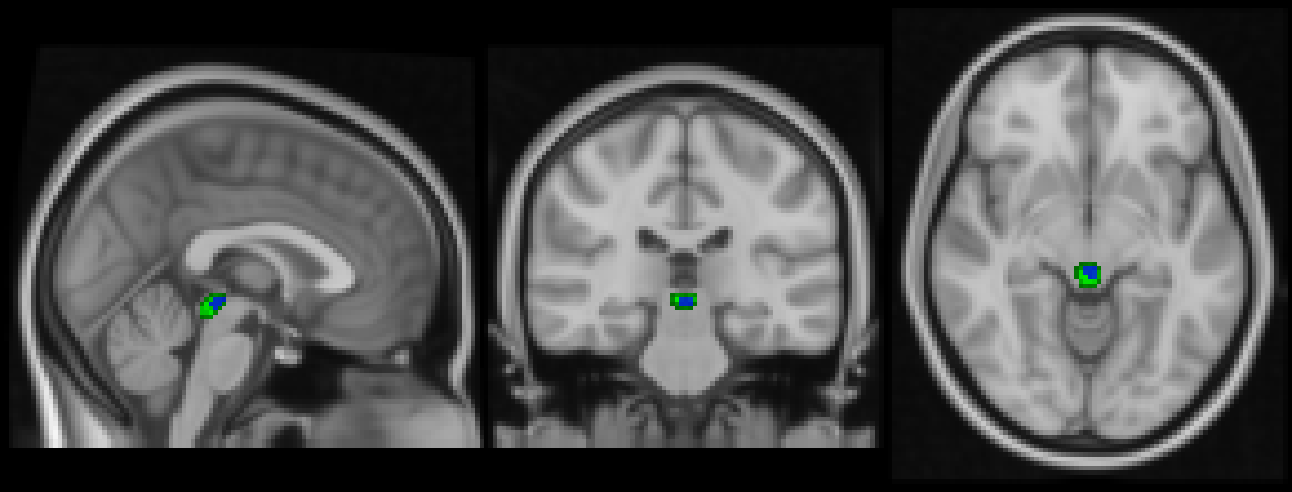

**Fig. S1 Periaqueductal gray seed.** The structural boundaries of the periaqueductal gray (PAG) were traced using the Duvernoy Atlas (in green). The overlap between the functional localizer and the structural seed was used in our analysis (in blue).

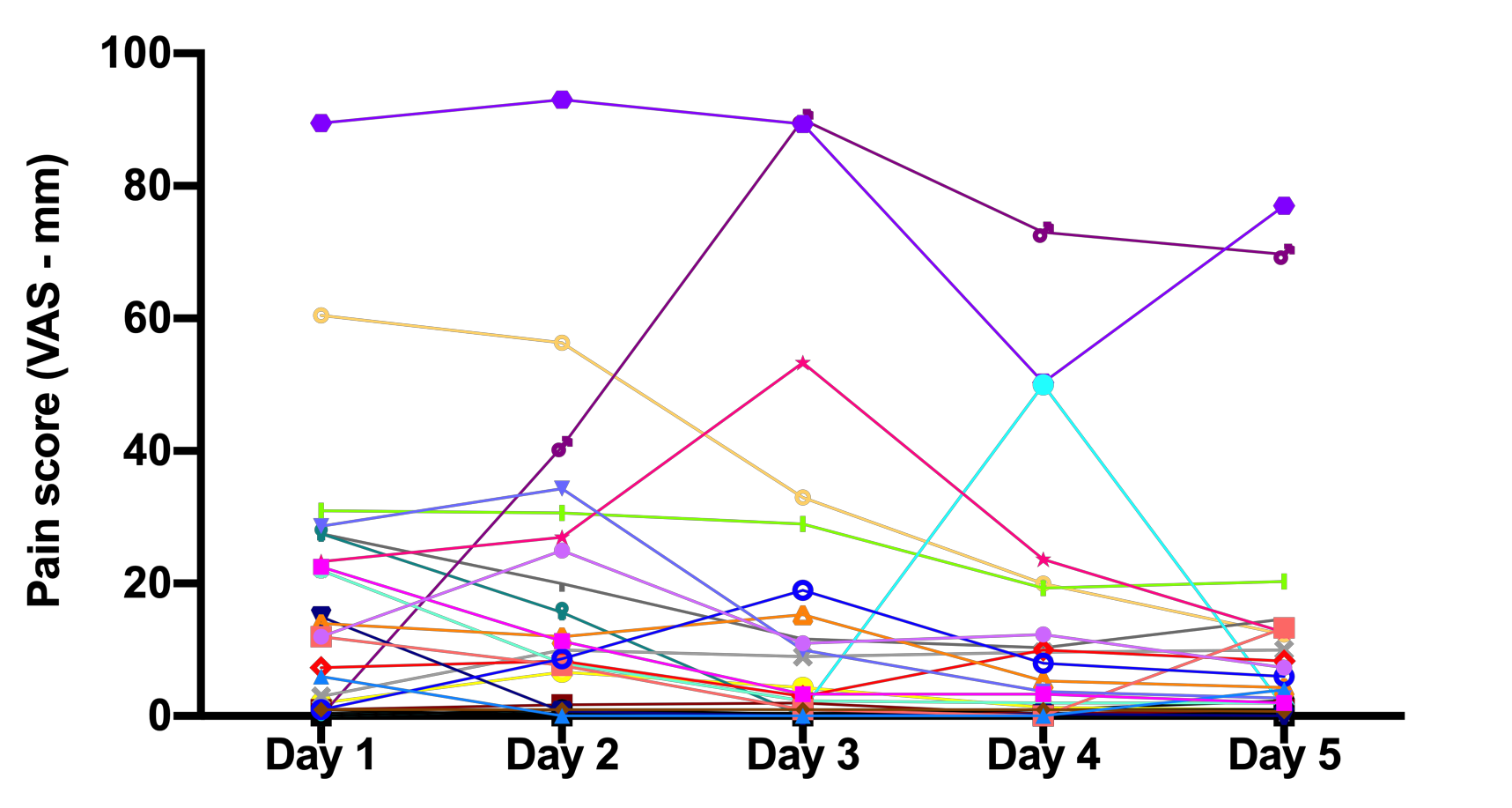

**Fig. S2. Individual daily pain ratings** Self-report pain ratings are represented following the insertion of the orthodontic separator over the course of five days for each participant (n=26). Participants reported their pain three times daily on a 100-mm visual analogue scale (VAS), and the average of these three ratings was used as a daily pain rating.
